# Uncovering genomic regions controlling root quality traits in Cassava (Manihot esculenta Crantz) using different GWAS models

**DOI:** 10.64898/2026.06.11.731598

**Authors:** Diana C Solarte Certuche, Gabriel M de Freitas, Jean-Luc Jannink, Cinara Fernanda Garcia Morales, Tamires Sousa Cerqueira, Bruna Santos de Santana, Eder Jorge de Oliveira, Antônio Augusto F Garcia

## Abstract

Cassava is a major staple crop in tropical regions, and improving its root nutritional quality, particularly carotenoid and dry matter content (DMC), remains a central breeding goal. To elucidate the genetic basis of these traits by locating genomic regions associated with them, we analyzed 3,043 cassava clones from the Brazilian Agricultural Research Corporation (Embrapa) breeding program, phenotyped across 188 multi-environment trials conducted from 2011 to 2022 in Brazil. All clones were genotyped using Genotyping-by-Sequencing (27,045 Single Nucleotide Polymorphism - SNPs) and Diversity Arrays Technology - DArTseq (25,923 SNPs). Trait values were estimated using a two-stage mixed model to obtain deregressed BLUPs (Best Linear Unbiased Predictions), and genome-wide association analyses were performed using both the Mixed Linear Model (MLM) and Multi-Locus Mixed Model (MLMM). We detected six significant SNPs consistently associated with carotenoid content and DMC after Bonferroni correction. These SNPs mapped to six candidate genes involved in pathways relevant to root physiology, including Abscisic Acid ABA-related signaling, hydrolase activity affecting carotenoid conversion, fatty-acid biosynthesis within plastids, cell-wall remodeling, and glycolytic energy metabolism. The loci jointly explained 75.56 % of the phenotypic variance for carotenoids and 76.23 % for DMC, with individual SNP effects ranging from ∼17 % to ∼42 % PVE (Proportion of Variance Explained). Broad-sense heritability was H2 = 0.78 for carotenoids and H² = 0.34 for DMC, confirming substantial genetic control and suitability for molecular breeding. Haplotype analyses revealed four superior haplotypes for carotenoids and one key haplotype for DMC, each showing significantly higher trait values compared with other allelic combinations. These haplotypes represent promising targets for marker-assisted selection and genomic selection, with direct applicability for accelerating genetic gain in elite breeding populations. The results provide actionable genomic resources for breeding programs aiming to develop biofortified and high-root quality cultivars and establish a foundation for future multi-omics and functional validation studies.

## Introduction

Cassava is recognized as the fourth most important staple food worldwide, providing carbohydrates, calories, calcium, and vitamins A, B and C (Kim & Iida, 2023). It serves as a dietary foundation for approximately 1 billion people across different continents. This adaptable crop demonstrates resilience to climate variability and the ability to thrive in marginal soils, making it a significant contributor to food security, industry, and economic development, particularly for rural communities (Forkum et al., 2025). Approximately 50 % of the world’s cassava production is concentrated in Africa, while Asia accounts for 38%, and Brazil ranks fifth with 5 % of the global output (Borku et al., 2025). Fresh cassava roots contain a high-water content of 70 %, along with 24 % starch, 2 % fiber, and 1 % protein. Other minerals such as iron, zinc, and provitamin A make up another 3 % of the roots. Starch is the predominant metabolite in cassava roots, accounting for 85 % of their dry weight, and it serves as a valuable raw material for human consumption, animal products, and industries producing biofuels and other derived products (Forkum et al., 2025).

Cassava is a great source of carbohydrates, but it has a low content of micronutrients. In human nutrition, vitamin A is crucial for vision and for the synthesis and differentiation of epithelial cells (Forkum et al., 2025). In developing countries, cassava is a staple food, making it difficult for people to meet their vitamin A requirements, particularly for pregnant women and children. This has become a public health issue (Qi et al., 2025). Increasing the carotenoid content in cassava roots is a key strategy for enhancing the biofortification of cassava, which aligns with the goals of both public health initiatives in developing countries and breeding programs (Saba et al., 2024). Carotenoid synthesis in plants is regulated by various hormones, such as jasmonic acid, salicylic acid, and abscisic acid (ABA). These hormones increase the accumulation of these pigments in the roots. Additionally, plastid division plays a significant role in the synthesis and accumulation of carotenoids (Sun et al., 2022). Respectively, with dry matter content, this is more strongly related to starch accumulation; the physiological processes that are important parts are the enzymes involved in the conversion of glucose, protein, and starch synthesis through sucrose synthase and cell wall-bound enzymes (Jia et al., 2025).

Considerable efforts have been made to develop cassava cultivars with yellow-orange flesh that are rich in β-carotene and provitamin A, as well as to enhance the dry matter content in elite cultivars. However, achieving genetic gains in these traits is challenging due to several factors, including high environmental variability, negative correlations between traits, polygenic inheritance, and complex physiological pathways. This highlights the necessity of complementing traditional breeding methods with genomic-assisted breeding tools to expedite the selection process and create cultivars that exhibit high dry matter content and elevated carotenoid levels. Recent studies in cassava, including research in proteomics and metabolomics (Olayide et al., 2023), transcriptomics (Xiao et al., 2021), and genomics of African cultivars (Villwock et al., 2025) (Rabbi et al., 2017), as well as investigations into phenotypic diversity in Brazilian germplasm (de Carvalho et al., 2022), have provided the cassava community with a deeper understanding of the genetic foundations of these traits. Furthermore, these studies elucidate the interactions among different metabolic pathways that influence the enhancement of these traits in specific cassava populations. Currently, there are no efficient markers for the Brazilian population that can be applied in routine breeding programs, and most of the GWAS (Genome-Wide Association Studies) studies made so far in cassava research for these traits has been used African germplasm. While these studies have identified significant regions on chromosomes one and ten, limitations regarding population structure have hindered the application of these markers within Brazilian germplasm. The integration of high carotenoid content and dry matter into breeding pipelines entails considerable costs and time investments associated with morphological evaluations and methods for determining dry matter content. Furthermore, the chemical procedures required to assess carotenoid content impose significant financial burdens. This situation presents an opportunity to analyze this information and select traits for molecular breeding, with the aim of enhancing carotenoid content in cassava roots. By pursuing this strategy, it is possible to accelerate processes, reduce costs, and improve efficiency within breeding program pipelines through the application of molecular markers.

GWAS leverages various natural and breeding populations, eliminating the need for a segregating population as seen in linkage mapping. GWAS can simultaneously detect multi-allelic variations and rare, small genetic effects, providing high-resolution insights into historical recombination events within the studied populations (Cantor et al., 2010). These models incorporate population structure through principal components and kinship matrices, along with covariates (Yu et al., 2006). This helps reduce false positives and allows the detection of major quantitative SNPs (single nucleotide polymorphisms) that might explain a significant portion of the phenotypic variation of traits (Briollais et al., 2016). Combining different GWAS models with haplotype-based analysis boosts the power of SNP detection and candidate gene identification.

In this study, we systematically investigated the genetic architecture of dry matter and carotenoids in cassava by locating genomic regions associated with them. We analyzed a population of 3,043 cassava clones, using a historical dataset to evaluate both dry matter content and carotenoid content. A GWAS was conducted using 27,045 SNPs from genotyping by sequencing and 26,000 SNPs from Diversity Arrays Technology datasets. The objectives of this study were as follows: (1) to characterize the phenotypic variation and population structure of the sample; (2) to determine genetic parameters, such as linkage disequilibrium, heritability, and correlations between traits; (3) to implement GWAS and haplotype analysis to gain insights into the genetic architecture of carotenoids and dry matter within the panel; and (4) to identify significant polymorphisms associated with candidate genes and superior haplotypes. The findings offer valuable tools for accelerating the breeding process aimed at biofortification and enhanced productivity in cassava roots, while also contributing to our understanding of the genetic architecture of these traits.

## Material and methods

### Panel of accessions and genotyping

The association panel comprised 3,043 clones from the germplasm of the Embrapa cassava breeding program, some of which were derived from various genomic selection cycles. Detailed information about this panel can be found in (https://www.nextgencassava.org/). DNA extraction for the cassava clones was performed using the CTAB method by Doyle & Doyle, 1987, and a protocol recommended by Diversity Arrays Technology Pty Ltd, Australia (Sansaloni et al., 2011). The clones were genotyped using two approaches: Genotyping by Sequencing (GBS) and Diversity Arrays Technology (DArTseq) SNP filtering, read alignment to the cassava reference genome (v6), and variant calling were performed using the *Genome Analysis Toolkit (GATK)* pipeline (McKenna et al., 2010). Standard quality control criteria were applied, excluding markers with a minor allele frequency (MAF) below 0.01 and loci with more than 60 % missing genotype data. This process yielded a total of 27,045 SNPs from GBS and 25,923 SNPs from DArTseq. These high-quality SNPs, which underwent further filtering, were utilized for the analysis presented in this study.

### Field experiments and Phenotyping data collection

Field experiments and phenotypic data collection were conducted across 188 trials performed over multiple years and locations in the state of Bahia, Brazil (Supplementary Table 1). The evaluations included various germplasm and genomic selection populations (C0, C1, C2, and C3), assessed using incomplete block, augmented block, and randomized complete block designs, depending on the year and location. Plot management, planting density, and standard crop husbandry followed Embrapa’s recommended protocols for cassava (Bernardo, 2020). Dry matter content was determined using the method described by Kawano et al., (1987), which is based on the linear relationship between dry matter (DM) and starch content, expressed as: DM = 158.3x – 142, where x represents specific gravity. Specific gravity was obtained as follows: (1) collect approximately 3–5 kg of roots and clean them to remove soil and debris; (2) weight the sample in air (*Wa*); (3) weight the same sample submerged in water (*Ww*) using the same container; (4) calculate specific gravity as: 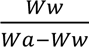; and (5) compute dry matter content using the equation DM = 158.3x – 142.

Carotenoid content was assessed according to the protocol described by Fukuda et al. (2010), using a visual color chart to score the intensity of yellow pigmentation in the parenchyma. Colors were classified on a five-point scale: 1 = white, 2 = cream, 3 = yellow, 4 = orange, and 5 = pink.

### Population structure and linkage disequilibrium analysis

Population structure and linkage disequilibrium analyses were performed to assess genetic relatedness among 3,043 clones using 27,045 SNPs generated by GBS and 25,923 SNPs generated by DArTseq. Population structure was inferred through Bayesian clustering in *STRUCTURE version 2.3.4* (Pritchard et al., 2000), with independent runs for K values from 1 to 6. The most likely number of clusters (ΔK) was identified using *STRUCTURE HARVESTER v.0.6.9.94* (Earl & vonHoldt, 2012), based on the log-likelihood of the data [LnP(K)] and the method of Evanno et al. (2005). Principal component analysis (PCA) was conducted in *TASSEL software v. 5.0* (Bradbury et al., 2007) to detect the subpopulation structure within the panel. Additionally, we used TASSEL to construct a neighbor-joining tree through phylogenetic analysis to explore the genetic relationships among the clones. The linkage disequilibrium (LD) was measured as the squared Pearson correlation between SNPs as r^2^ = (cor(*X*_i_, *X*_j_))^2^ where *X*_i_ and *X*_j_ are genotypes vector of the two SNPs (Hill & Robertson, 1968)(Rogers & Huff, 2009) this calculation was doing in population-base linkage tool (PLINK) (Purcell et al., 2007). Patterns of LD decay and local r² distributions were examined to help delineate candidate genomic regions. All visualizations were produced using standard plotting functions in R. the Kinship matrix was calculated using the VanRaden Method with the equation 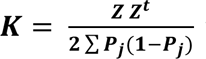 where k is the kinship matrix, **Z** is the genotype matrix and *P*_j_ is the frecuency of the allele in the SNP *j* (VanRaden, 2008).

### Phenotypic data analysis

Data was analyzed for each trait and across locations and year combination using a two-stage mixed-model framework adapted for multi-environment trial data using lme4 v.1.1-31 for R statistical software (Bates et al., 2015). In the model all factors were trated as random effects in the analyses, except the genotype effect to estimates Best Linear Unbiased Estimator (BLUEs, These blues were obtained using the first step in which environment-specific linear mixed models were fitted to plot-level of dry matter and carotenoids observations, adjusting for the design effects and obtaining genotype adjusted means (BLUEs/BLUPs) Best Linear Unbiased Prediction for each environment.

The model for this step is as follow:

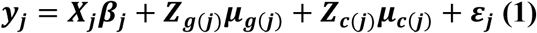

where, ***y*_j_** represents the data vector of PPD evaluation plot in the location-year combination that represents the environment. ***X***_***j***_is an incidence matrix relating observations in enviroment j, ***β***_***j***_ is a vector that represents the fixed effects specfici for environment *j^th^*, ***Z***_***g***(***j***)_is an incidence matrix for clones, ***μ***_***g***(***j***)_is a vector that contains the random effects of the clone in envrionmet *j^th^*, ***Z***_***c***(***j***)_incidence matrix for additional random desgin-realted effects in evironment *j^th^*, such as row and columns, and blocks, ***μ***_***c***(***j***)_vector of random effects between environments in the combination location-year, and, ***ε***_***j***_vector of plot error associated with the environment *j^th^*. Assumed that ***ε***_***j***_ ∼ ***N***(***0***, ***g***_***j***_), ***μ***_***g***(***j***)_ ∼ ***N***(***0***, ***g***_***j***_), ***μ***_***c***(***j***)_ ∼ ***N***(***0***, ***c***_***j***_).

In the second stage, environment-level genotype estimates were combined to obtain single genotype genetic values. De-regressed BLUPs (or weighted DBLUPs) were computed to account for heterogeneous error variances among environments. The second-stage working model can be expressed as:

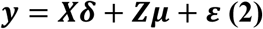

where ***y*** is the vector of de-regressed genotype values, ***X*** is the design matrix relating genotype level observations to fixed effects across environments, ***δ*** contains fixed effects (environmental, markers and inbreeding), ***Z*** is the incidence matrix assigning genotype-level observation to genotypes, ***μ*** denotes random genetic effects structured by marker-based relationships, and ***ε*** is the residual vector.

The genetic value of each individual estimated as BLUP from the mixed linear models is process by the procedure describe by (Garrick et al., 2009). The BLUPs were de-regressed using the following formula:

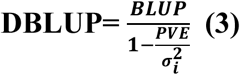

Where *PEV* represented the prediction error variance for each BLUPs and 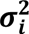 is the clonal variance. This procedure allows to avoid applying shrinkage to the same data. This DBLUPs were used in the genomic prediction analyses.

The resulting DBLUPs values obtained in this analysis were using to the GWAS analysis model in the next steps.

Broad sense heritability for the trait was estimated on an entry mean basis using the equation

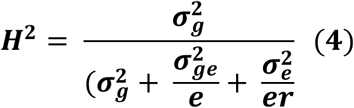

Where 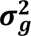 is the genotypic variance, 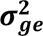 is the genotype by environment interactions variance, 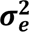 error variance, and respectively **r** and e the number of replications and enviroments within each of the environments included in the analysis.

### Genome-wide association study of root quality traits

A GWAS analysis was conducted using 27,045 SNPs from GBS and 25,923 SNPs from DArTseq across 3,043 clones with DBLUP values for DMC and carotenoid content. Different statistical models were employed to enhance power and reduce false positives, including the Multi-locus Mixed Model (MLMM), which adjusts for cofactors associated with markers through forward and backward stepwise regression in a mixed model. The Bayesian Information Criterion (BIC) is used to determine the optimal number of cofactors. In the forward step, SNPs are added as cofactors to prove if they improve the model. Taking a backward step removes cofactors that are not significant (Segura et al., 2012). Additionally, False Discovery Rate (FDR) was utilized to improve the control of Type I and Type II errors, proving effective with family-based and structured association samples (Yu et al., 2006). Both models were implemented in Genome Association and Prediction Integrated Tool (GAPIT) version 3 R package (J. Wang & Zhang, 2021) taking into account both the kinship K (VanRaden, 2008) and population structure Q matrices, For multiple testing correction, the Bonferroni procedure was applied with a significance level of α = 0.05 was used to determine the threshold p-value (P=0.05/n, where n = number of SNP), the markers were considered significant when p-value was less than 1.848 x 10^-6^. The results were visualized using Manhattan plots, with a nominal threshold set at −log10(p-value) = 5.5, and quantile-quantile (Q-Q) plots were created to evaluate statistical inflation and model fit. Linkage disequilibrium (LD) was estimated through pairwise correlation coefficients (r²) to identify candidate regions, and loci with −log10(p) > 3.6 (p < 2.51 × 10⁻⁴) were selected for further investigation. The PVE (Phenotypic variance explained) were calculated using the formula 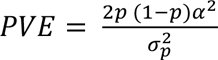 where 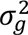 is the genetic variance and p is the allele frequency, *α*^2^is the estimate effect of the SNP and 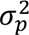 is the total phenotypic variance (Zhang et al., 2010).

### Candidate gene annotation and haplotype block analysis

The decay of LD estimated at a physical distance was 1 kb for the genotypic dataset, this information was used as the reference window for identifying candidate gene identification and haplotype analysis. Genes related to dry matter content and carotenoids within ±1 kb of significant SNPs were annotated using the cassava reference genome v6.1. (Bredeson et al., 2016). Functional information was retrieved from *Gene Ontology (GO)* (https://geneontology.org/), *InterPro* (https://www.ebi.ac.uk/interpro/) and additional annotations available through *National Centre for Biotechnology Information* (NCBI) (http://www.ncbi.nlm.nih.gov/) and *European Molecular Biology Laboratory European Bioinformatics Institute* (EMBL-EBI) (https://www.ebi.ac.uk/). When relevant, gene annotations were cross-referenced with previously published studies addressing the traits analyzed in this work. Haplotype block analysis was performed using *HaploView v.4.2* (Barrett et al., 2005), and gene plots were generated with *SnapGene v.7.0* (www.snapgene.com). The haplotype blocks were defined based on the LD decay results from the GWAS analysis.

## Results

### Phenotypic variation, descriptive statistics and correlation

We investigated phenotypic variation in a dataset that included dry matter content and carotenoid content, evaluated using a visual scale. The dry matter content in the studied population ranged from 16.11 to 50.98 %, with a mean of 34.65 %. Most clones displayed a rating of 1 (79.29 %), while a smaller proportion received ratings of 2 (13.94 %), 3 (6.57 %), and 4 (0.1 %) for carotenoid content. Overall, the population ranged between ratings of 1 and 4. The distribution of dry matter content followed a normal distribution, with most clones having values between 30 and 37.5 % (Supplementary Figure S1.

The broad-sense heritability for carotenoids was estimated at H² = 0.78, indicating a high genetic contribution to phenotypic variance in this trait. In contrast, the broad-sense heritability for dry matter was H² = 0.34, indicating a moderate level in cassava. The genotype-by-environment variance was recorded as 0.43 for carotenoids and 0.45 for dry matter, suggesting similar variances for both traits,. Pearson’s correlation analysis revealed a negative correlation between dry matter content and carotenoid DBLUPs, with a coefficient (r = −0.10) and a p-value < 0.05 (Supplementary Figure S2).

### Marker distribution, population structure and kinship

A total of 27,045 SNPs from GBS and 25,923 SNPs from DArTseq were mapped across the cassava genome, covering an estimated 35 Mb (Supplementary Figure. S3). In both datasets, chromosome 1 contained the highest number of SNPs (n = 572), whereas chromosome 14 contained the fewest (n = 259) (Supplementary Figure. S3A, S3C). The distribution of minor allele frequencies (MAF = 0.05–0.50) showed an average MAF of 0.24 (Supplementary Figure. S3B, S3E), and few markers exhibited allele frequencies below 5 %, providing sufficient genomic resolution for downstream association analyses.

The population structure of the 3,043 cassava clones was assessed using structure inference. Principal component analysis (PCA) revealed two major stratifications within the panel. The first two principal components (PC1 and PC2) together accounted for 9.39 % of the total genetic variance, and the inclusion of PC3 increased the cumulative variance explained to approximately 11.72 % (Figures. 1A–B). Bayesian clustering, informed by kinship estimates, also supported the presence of two genetically distinct groups within the population (Figure. 1C). One cluster corresponded to historical Embrapa germplasm, whereas the second cluster comprised clones derived from genomic selection cycles. Identity-by-state estimates and the kinship heatmap indicated generally low pairwise relatedness among most samples, consistent with the presence of broad genetic diversity within the panel.

**Figure 1.**
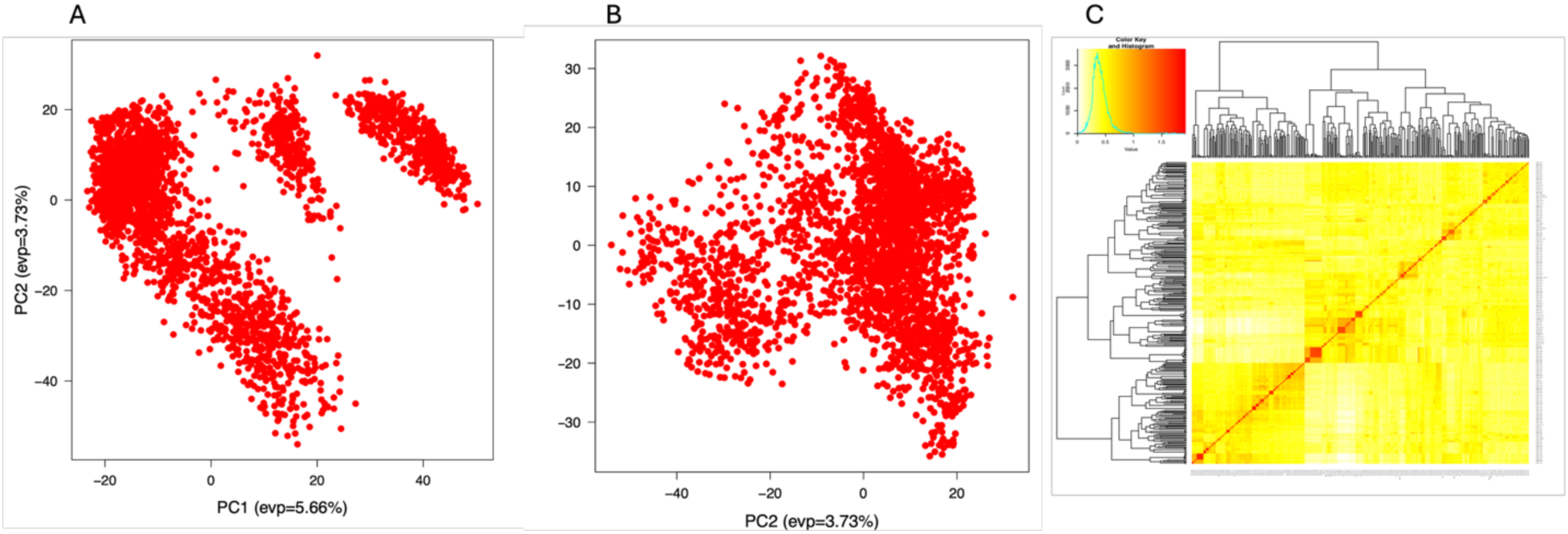
Population structure assessed through principal component analysis (PCA). **A)** Scatterplot of the first three principal components (PC1–PC2) showing sample distribution across major axes of genetic variation. **B)** Scatterplot of the first three principal components (PC2–PC3) showing sample distribution across major axes of genetic variation **C)** Heatmap of the pairwise kinship matrix, where yellow and red represent weak and strong relatedness, respectively. The hierarchical clustering dendrogram is shown along the matrix margins.

### Genome-wide association analysis and linkage disequilibrium

In the DArTseq dataset, LD started at r² = 0.58 and decayed to approximately r² ≈ 0.12, whereas in the GBS dataset, the initial r² was 0.40, decaying to about r² ≈ 0.18 (Supplementary Figure. S3C and S3F). Based on this pattern, a window of ∼1 kb around significant SNPs was selected for defining candidate genomic regions.

Using the MLMM and MLM models in the GWAS analyses, we identified a total of two SNPs significantly associated with dry matter content and five SNPs associated with carotenoids (Supplementary Table S1). The Q–Q plots (Figures. 2A and 2B) showed only a slight deviation from the expected distribution for both traits, indicating appropriate control of population structure and relatedness. The Manhattan plots revealed multiple SNPs that exceeded both the Bonferroni-adjusted significance threshold of −log10(p) = 5.5 For carotenoids, four SNPs were identified on chromosome 1 by the MLMM model, while the MLM model recovered two SNPs on the same chromosome. Two of these loci were consistently detected by both models. Additional carotenoid-associated SNPs were identified on chromosomes 4 and 10, with each model uniquely identifying one SNP. For dry matter content, two SNPs were identified by both MLMM and MLM: one on chromosome 1 and one on chromosome 10 (Supplementary Table S1).

**Figure 2.**
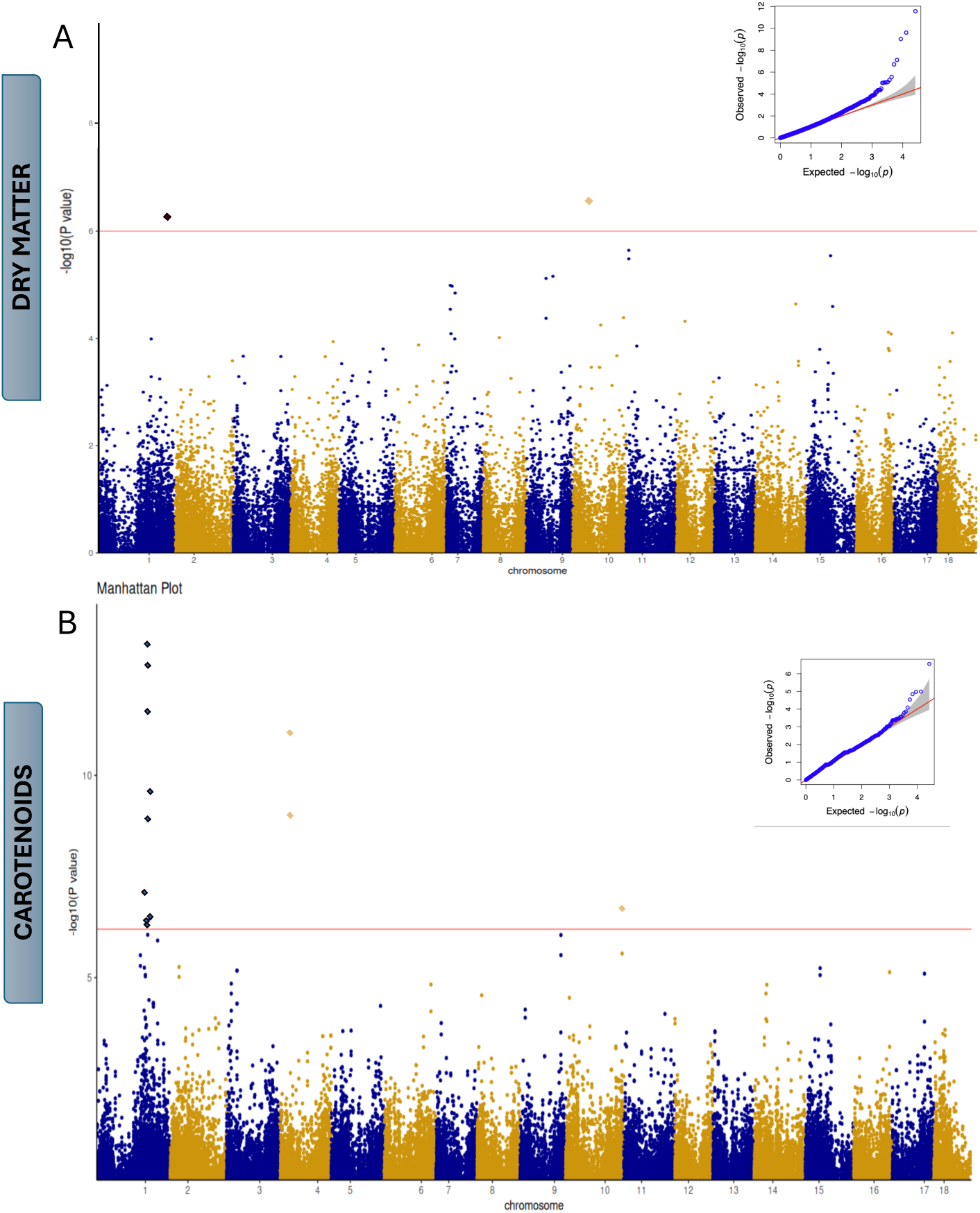
Results of genome-wide association studies (GWAS) using MLMM and MLM statistical models. **(A)** Manhattan plot and quantile–quantile (Q–Q) plot for dry matter content; **(B)** Manhattan plot and quantile–quantile (Q–Q) plot for carotenoid content.

On chromosome 1, the SNPs associated with carotenoid content, S01_22503611 (T/A), S01_23979768 (G/T), and S01_25281512 (T/C), collectively explained 42 % of the phenotypic variance (PVE). The SNP on chromosome 4, S04_5593052 (A/G), accounted for 32 % of the PVE, and the SNP on chromosome 10, S10_25598238 (T/C), accounted for 17.71 %. Together, these five SNPs explained 89.24 % of the phenotypic variance for carotenoid accumulation in the evaluated population. For dry matter content, the SNP on chromosome 1, S01_31462045 (T/C), accounted for 42.20 % of the PVE, while the SNP on chromosome 10, S10_7304197 (T/C), explained 34.02 %; jointly, these variants accounted for 76.23% of the total PVE. Comprehensive details regarding the peak SNPs associated with all loci and their candidate genes are provided in the Supplementary Table S1.

### Candidate genes associated with significant SNPs

Candidate gene annotation was performed for all significant SNPs identified in the GWAS. Seven genes were identified within the ±1 kb LD-based genomic windows surrounding peak SNPs, using the cassava reference genome in *Phytozome v6.1* (Goodstein et al., 2012). Two candidate genes were associated with dry matter content and five with carotenoid accumulation. For dry matter content, two SNPs were located in regions containing annotated genes. The SNP S01_31462045, on chromosome 1, was located within the coding region of *Manes.01G231200*, a gene annotated with functions related to cell wall modification (Figure 3A). The second SNP, S10_7304197 on chromosome 10, was positioned within a non-coding region adjacent to *Manes.10G055500*, which is annotated with iron ion–binding activity (Figure. 3B).

**Figure 3.**
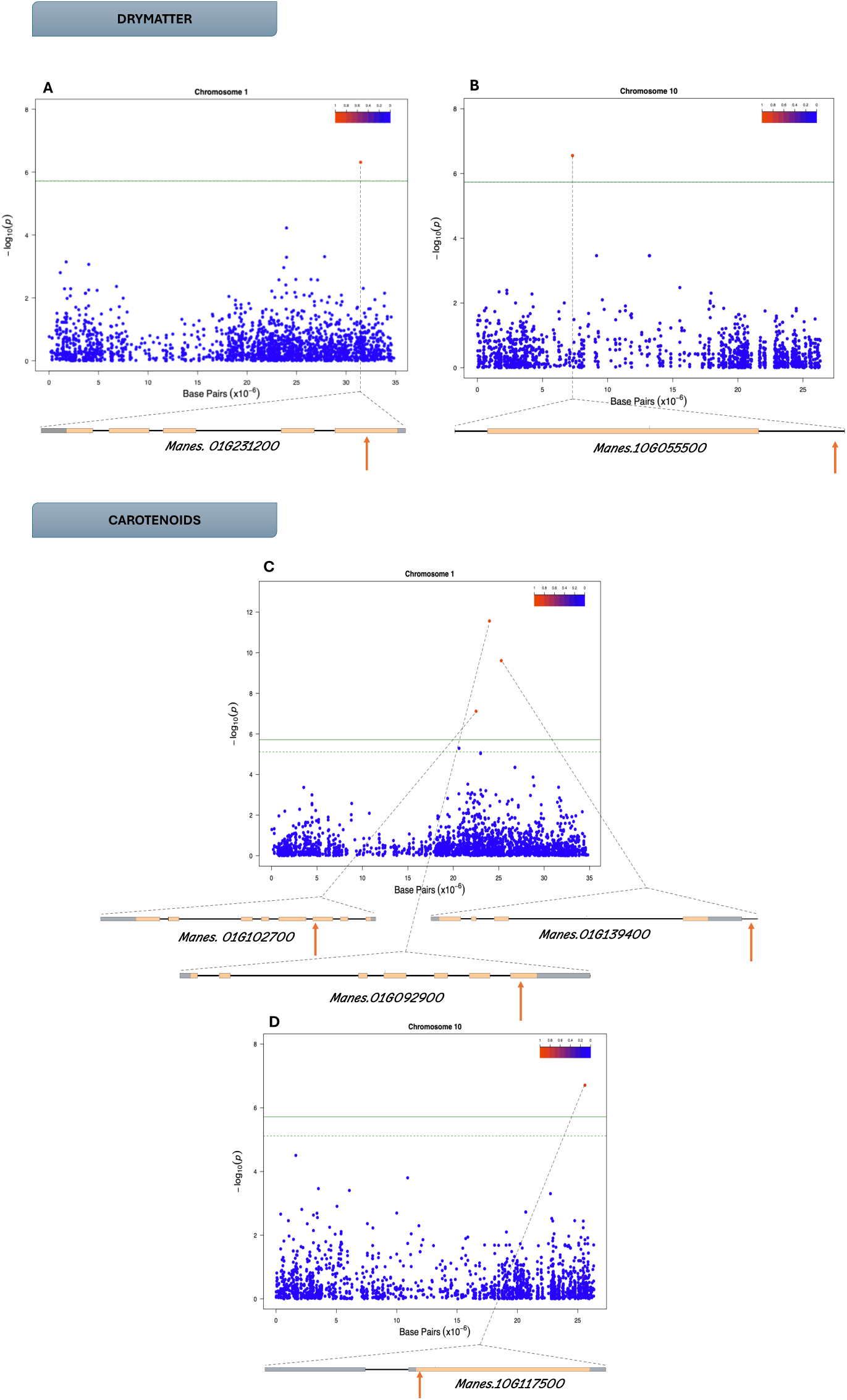
Candidate genes and local linkage disequilibrium (LD) around peak SNPs. (A–D) Regional association plots for dry matter content **(A, B)** and carotenoids **(C, D)** on chromosomes 1 and 10. The upper panels show the GWAS signals, and the lower panels present pairwise LD (r²; color scale) together with annotated gene models within the interval. The lead SNP for each region is indicated by a dashed line, and the genome-wide significance threshold is marked. Orange pointers denote SNPs mapped to gene models. coding regions are shown in gray, and coding sequences in bisque.

For carotenoids, five SNPs were associated with four annotated genes. On chromosome 1, the SNP S01_22503611 was located within the coding region of *Manes.01G102700*, annotated as a phosphatidylinositol-specific phospholipase C (Figure 3C). The SNP S01_23979768 was found within the coding region of *Manes.01G092900*, annotated with protein kinase activity. The SNP S01_25281512 mapped to a non-coding region adjacent to *Manes.01G139400*, annotated with hydrolase activity. On chromosome 10, the SNP S10_25598238 occurred within the coding region of *Manes.10G117500*, annotated as D-3-hydroxyoctanoyl-[acyl carrier protein] dehydratase, an enzyme associated with fatty acid biosynthesis (Figure 3D). On chromosome 10, a single SNP (S10_25598238) was identified within the coding region of *Manes.10G117500*. This gene is annotated as encoding D-3-hydroxyoctanoyl-[acyl carrier protein] dehydratase, an enzyme associated with the fatty acid biosynthetic process (Figure 3D).

### Haplotype analysis

Haplotype structure was examined for genomic regions flanking the significant SNPs on chromosomes 1 and 10 for both traits. For carotenoids, haplotypes were defined for the four genes identified by GWAS. On chromosome 1, the SNP S01_22503611 mapped to the coding region of *Manes.01G102700*, and the haplotype configuration showed a nonsynonymous substitution (Figure 4A). The SNP S01_25281512, associated with *Manes.01G139400*, was located in a non-coding region (Figure 4B). The SNP S01_23979768, located within *Manes.01G092900*, also resulted in a nonsynonymous substitution (Figure 4C). On chromosome 10, the SNP S10_25598238 mapped to *Manes.10G117500*, producing a synonymous substitution (Figure 4D). The haplotype frequencies and the distribution of allelic combinations across the population are shown in Figure 4E. Boxplots summarizing carotenoid accumulation for individuals carrying 0, 1 or 2 copies of the associated alleles are presented in Figure 4F.

**Figure 4.**
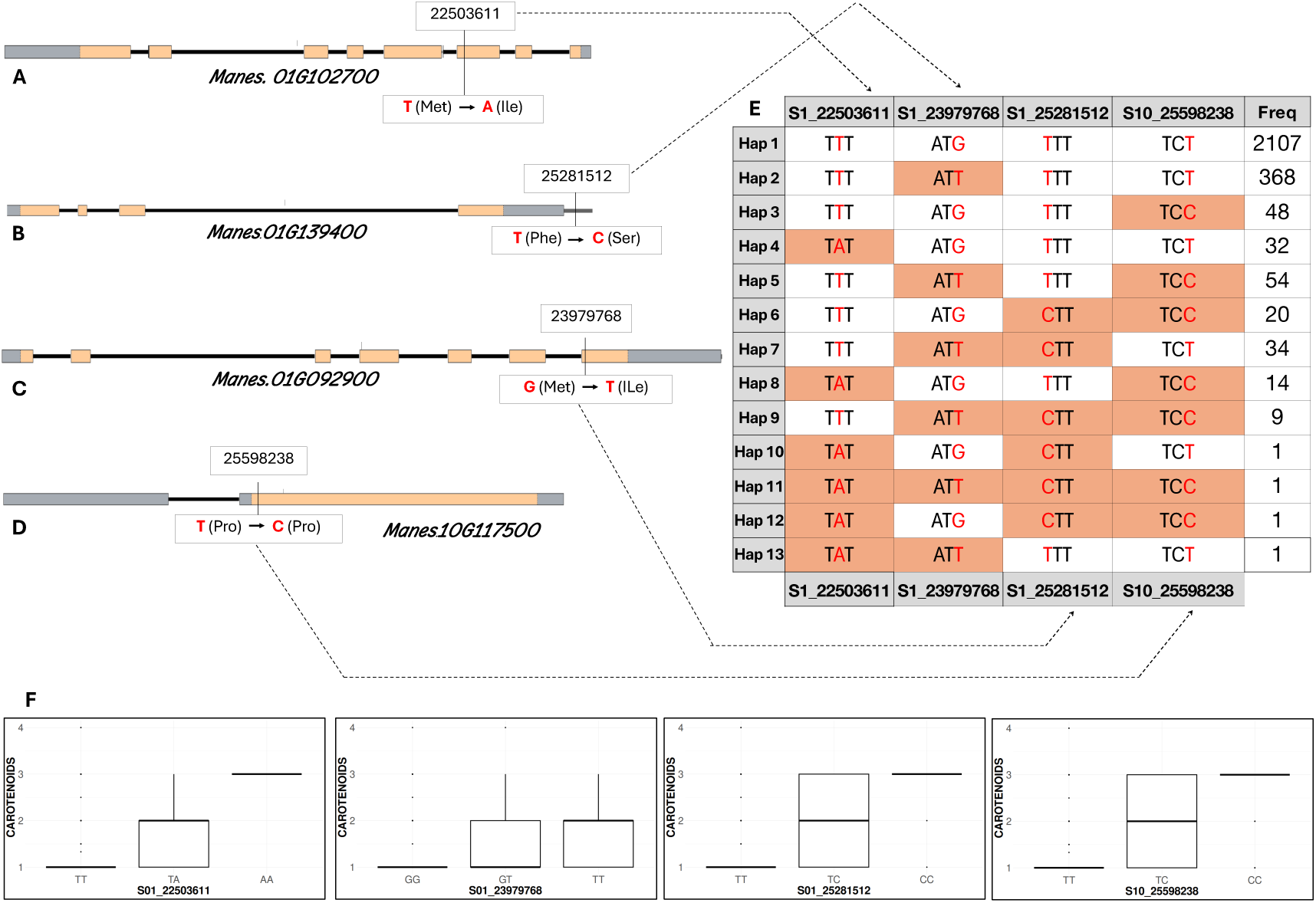
Haplotype analysis of candidate genes associated with carotenoid accumulation in cassava. **(A–D)** Haplotype maps for the candidate genes *Manes.01G102700, Manes.01G139400*, *Manes.01G092900*, and *Manes.10G117500*. Coding regions are shown in gray, coding sequences in bisque, mutated bases are highlighted in red, and amino-acid substitutions (where present) are indicated in parentheses. **(E)** Summary of SNP information for chromosomes 1 and 10, showing variant positions (gray), allele-combination frequencies in the population (right), number of haplotypes detected (left), and codon-level allele configurations (center). Codons are colored according to the favorable allele for the trait, with reference and alternative alleles highlighted in red. **(F)** Boxplots displaying carotenoid content for individuals carrying 0, 1, or 2 copies of the associated SNP alleles.

For dry matter content, the allelic variation corresponding to the two genes identified by GWAS analysis*, Manes.01G231200* presents a nonsynonymous substitution from Leucine (Leu) to Lysine (Lys). *Manes.10G055500* presents a synonymous mutation for Leucina.We located two SNPs, one in the chromosome one region in 31.46 Mb and the other in the chromosome ten region located in 7.3Mb. Together, these two regions form 4 haplotypes in the population under study, listening with frequencies as follow; Hap 1[TT] 2527(94 %), Hap 2[TC] 48(1.78 %), Hap 3[CT] 100(3.72 %), Hap 4[CC] 11(0.40 %) (Figure 5c). Haplotype 3 has both copies of the favorable alleles for both genes related to dry matter content; this suggests that these clones present a high accumulation of dry matter. In Figures 5d and 5e, the box plots show the phenotypic behavior of clones with 1 or 2 copies of the SNPs. In the case of SNP S01_31462045, these clones exhibit a better dry matter content, and clones that have zero copies of the SNP S10_7304197 present a better behavior with respect to this trait. These results, together with the GWAS results, allow us to have a better understanding of the genetic architecture of carotenoids and dry matter, leading to the identification of a favorable allele combination on chromosomes 1 and 10 for both traits, advancing the introgression of this genetic background into the future breeding population.

**Figure 5.**
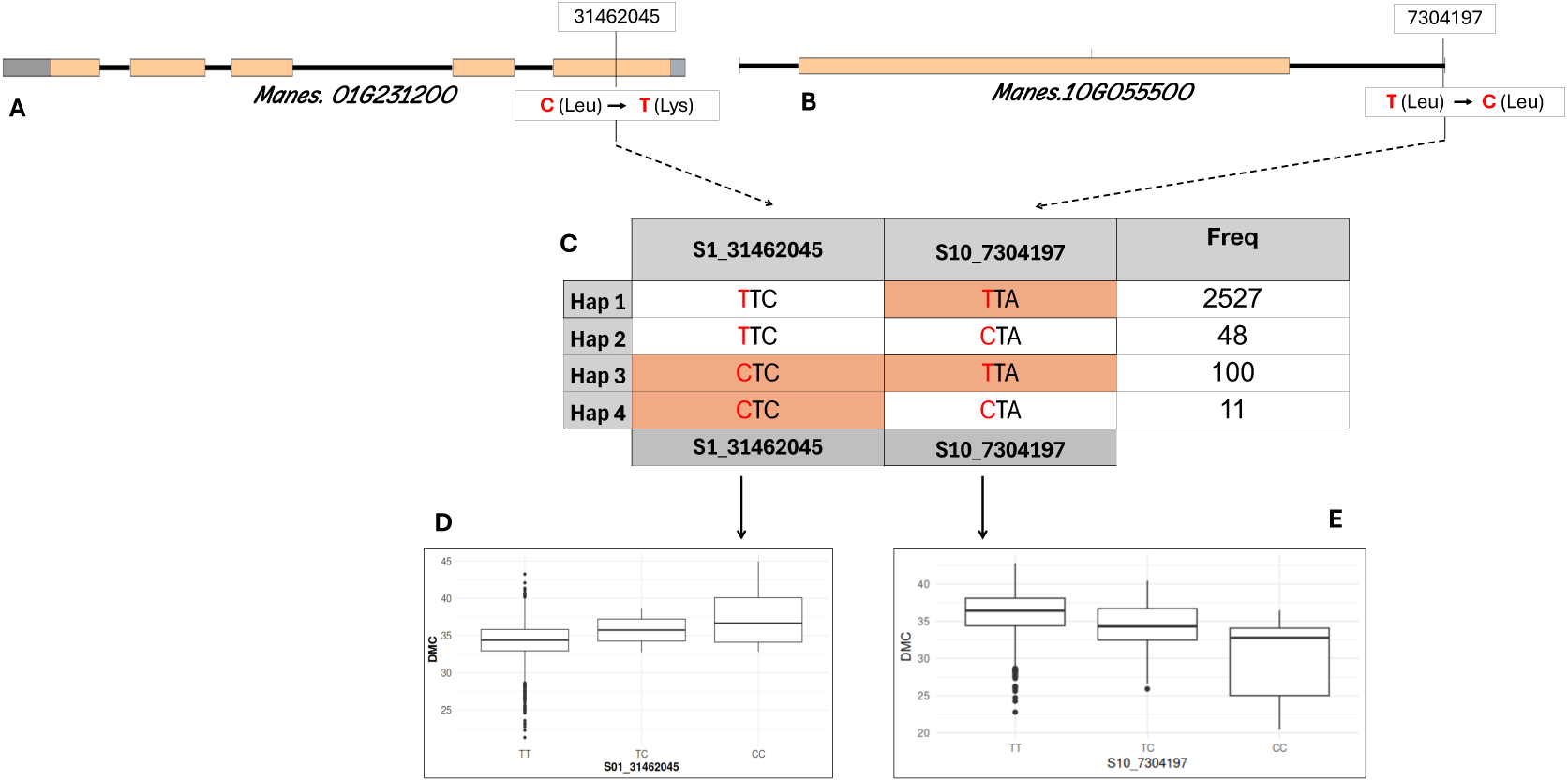
Haplotype analysis of candidate genes associated with dry matter content in cassava. **(A–B)** Haplotype maps for the candidate genes *Manes.01G231200* and *Manes.10G055500*, showing coding regions (gray), coding sequences (bisque), mutated bases (red), and amino-acid substitutions when present. **(C)** SNP summary for chromosomes 1 and 10, indicating variant positions (gray), allele-combination frequencies in the population (right), number of haplotypes identified (left), and codon-level allele configurations (center). Codons are colored according to the favorable allele for the trait, with reference and alternative bases highlighted in red. **(D–E)** Boxplots showing dry matter content for individuals carrying 0, 1, or 2 copies of the associated SNP alleles.

## Discussion

Dry matter and carotenoid content are key traits for root quality cassava, industrial processing, and biofortification initiatives (Saba et al., 2024). These two traits have certain limitations regarding their evaluation methods. For carotenoids, a visual scale is typically used because it is a quick and cost-effective method for analyzing a large number of samples (Jaramillo et al., 2018). However, this approach is subjective and can vary depending on the evaluator and the specific portion of the sample being assessed. Conversely, the specific gravity method utilized to assess dry matter lacks accuracy when applied across diverse genotypes, varying evaluation timelines post-harvest, and differing environmental conditions (Silva et al., 2023). Collectively, these limitations emphasize the need for a more comprehensive understanding of the genetic architecture that underlies these traits, particularly the genomic regions involved, the associated genes, and the allelic combinations that contribute to phenotypic expression. This is essential for designing more efficient selection strategies in cassava breeding. In this study, integrating multi-year phenotypic data with high-density SNP genotyping enabled the identification of loci, candidate genes, and haplotype patterns associated with DMC and carotenoids in a large breeding panel representative of Brazilian germplasm.

### Phenotypic variation, genetic variance, and trait correlations

Substantial phenotypic variation and high genotypic variance were observed for both traits, demonstrating that the population holds sufficient diversity for effective genome-wide association analyses. Dry matter content exhibited a wide quantitative range, whereas carotenoid levels, evaluated via visual score, showed narrower phenotypic dispersion but high genotypic signal. The moderate negative correlation between DMC and carotenoids (r = – 0.10) aligns with prior findings in African germplasm (Villwock et al., 2025), confirming that the relationship between starch accumulation and carotenoid biosynthesis is weak. Thus, simultaneous genetic improvement of both traits appears feasible, with minimal antagonistic pleiotropy. For breeding programs, this is an advantageous scenario, as it suggests that gene pyramiding for DMC and carotenoids can be pursued without substantial risk of unfavorable interactions between alleles influencing the two traits.

In the present study, we observed significant genotypic variance and a wide range of phenotypic variation for DMC. We found strong genetic variance for carotenoids associated with different alleles in the population. This indicates that the population possesses sufficient genetic diversity to generate robust GWAS results. These findings are in agreement with earlier studies involving African germplasm (Rabbi et al., 2017) and Brazilian germplasm (Sampaio et al., 2024), both of which similarly report substantial genetic variation within their evaluated populations.

Heritability estimates reinforce the potential for high genetic gain. Broad-sense heritability was moderate for DMC (H² = 0.34) and high for carotenoids content (H² = 0.78), reflecting a high genetic basis, particularly for carotenoids, and favorable prospects for selection. These values differ slightly from those described by Villwock et al. (2025), who reported H² = 0.49 for carotenoids and H² = 0.50 for DMC in African germplasm, but remain within the expected range for clonally propagated crops evaluated across multiple environments.

### Linkage disequilibrium, population structure, and GWAS performance

LD decay is a key parameter influencing the resolution and statistical power of genome-wide association studies (Briollais et al., 2016). LD patterns reflect underlying evolutionary and demographic forces—including recombination rates, selection, and population admixture, which collectively shape the extent to which alleles at different loci are inherited together (Cantor et al., 2010). In this study, LD decay occurred rapidly within ∼1 kb, with r² declining from 0.58 to background levels. This rapid decay indicates a high degree of historical recombination and substantial genetic diversity within the breeding population, consistent with repeated cycles of selection and recombination incorporated into Embrapa’s cassava breeding pipeline. Such LD patterns provide favorable conditions for fine-mapping loci and increase the precision of GWAS interpretation.

Population structure analysis revealed two major subgroups within the panel; one consisting of historical Embrapa germplasm and the other representing clones derived from genomic selection cycles. Accounting for this structure was essential for reducing spurious associations, as reflected by the well-aligned Q–Q plots and clean Manhattan plot profiles. The genetic distinction between these two groups suggests that both historical diversity and more recently recombined backgrounds contribute to the observed variation, reinforcing the value of this panel for association mapping.

The GWAS analysis results provide a robust number of SNPs detected across the genome for both traits, although with a high value of PVE. These SNPs result from the different models used, suggesting that each model and the genetic algorithms implemented allow better results and a larger number of SNPs detected from the datasets. The MLMM recovered a greater number of associations than MLM, while both models effectively controlled inflation and reduced false positives. For dry matter content, both MLMM and MLM consistently detected significant SNPs on chromosomes 1 and 10. For carotenoids, MLMM identified four SNPs across chromosomes 1, 4, and 10, whereas MLM detected three SNPs on the same chromosomes (Supplementary Table S1). These findings are consistent with earlier GWAS studies in African cassava germplasm (Rabbi et al., 2017; Villwock et al., 2025), which also reported significant associations on chromosomes 1 and 10 for both carotenoids and dry matter content. The convergence of results across diverse geographic backgrounds suggests that the genetic architecture of these traits may be conserved in global cassava populations.

### Candidate gene discovery

GWAS have become a powerful approach for identifying genomic regions and putative causal genes underlying key agronomic traits in cassava. In this study, GWAS proved highly effective for detecting loci associated with two physiologically complex traits, carotenoid and dry matter content, within a genetically diverse breeding population. Consistent with the observed rapid LD decay, the high recombination rate and diversity within the panel enabled high-resolution mapping. Overall, six significant SNPs and six candidate genes were identified as strongly associated with variation in carotenoids and dry matter content.

For carotenoids, four genes were associated with SNPs that together explained 75.56 % of the phenotypic variance. On chromosome 1, the SNP S01_22503611 mapped to *Manes.01G102700*, annotated as a phosphatidylinositol-specific phospholipase C. Enzymes in this class participate in lipid-associated signaling cascades and are connected to transcriptional regulation involving ABA and Ca²⁺—pathways broadly implicated in carotenoid biosynthesis and plastid development (Drapal et al., 2024). Elevated ABA signaling in high-carotenoid cassava genotypes was also reported by Olayide et al. (2023), further supporting the importance of this regulatory network. Another SNP on chromosome 1, S01_23979768, was located near *Manes.01G121900*, a gene annotated with protein kinase activity. Protein kinases are known regulators of ABA-mediated signaling processes and have been linked to metabolic adjustments associated with carotenoid accumulation (Bender & Zipfel, 2023). Together, these two genes work to activate and regulate ABA functions during the accumulation of carotenoids. This interaction is also documented by (Xiao et al., 2021), who emphasize the relationship between kinase activity and the ABA signaling pathway, which influences β-carotene accumulation in cassava roots.

The SNP S01_25281512 mapped to *Manes.01G139400*, annotated with hydrolase activity. Hydrolases, along with desaturases and cyclases, participate in enzymatic steps that influence flux through the carotenoid pathway, particularly those related to the conversion of lycopene to β-carotene, a relationship also emphasized by transcriptomic evidence in Olayide et al. (2023). On chromosome 10, SNP S10_25598238 was found within *Manes.10G144300*, which encodes D-3-hydroxyoctanoyl-[acyl carrier protein] dehydratase, a key enzyme in the fatty acid synthase II complex (Drapal et al., 2024). Fatty-acid biosynthesis is essential for the formation and maintenance of plastid membranes and lipid-rich structures, which serve as sites for carotenoid sequestration. Genes associated with this enzymatic complex were also reported to be highly expressed in high-carotenoid cassava genotypes (Olayide et al., 2023).

For dry matter content, two candidate genes were identified, collectively explaining 76.23 % of the phenotypic variation. The SNP S01_31462045 on chromosome 1 was located within *Manes.01G231200*, a gene involved in homogalacturonan degradation and cell wall modification. This gene has been previously linked to cell wall remodeling processes associated with starch accumulation and root development (Jia et al., 2025; Wang et al., 2016). The second SNP associated with dry matter, S10_7304197 on chromosome 10, mapped to *Manes.10G058150*, which encodes a protein annotated with iron ion binding and ATP/GTP binding activity. Such proteins participate in glycolytic pathways and energy mobilization and have been reported as highly expressed during tuberous root formation in cassava (Wang et al., 2016) supporting their role in processes associated with biomass and starch deposition. Together, these candidate genes represent biologically relevant pathways connected to carotenoid metabolism, plastid function, carbohydrate deposition, and root development. Their identification provides a strong foundation for future functional validation and incorporation into marker-assisted and genomic selection pipelines.

### Haplotype analysis and implications for cassava breeding

Haplotype analysis provided additional resolution into the allelic variation and functional polymorphisms underlying carotenoid accumulation and dry matter content in Brazilian cassava germplasm. For carotenoids, the gene *Manes.01G102700* exhibited a nonsynonymous substitution [T/C] that changes methionine to isoleucine. Individuals carrying the C allele showed higher carotenoid content (Figure 4F). Although the biochemical implications of this substitution require validation, isoleucine substitutions have previously been associated with altered enzyme activity in lipid-associated signaling pathways (Ohmura et al., 2001), which are known to influence carotenoid regulation. For *Manes.01G139400*, located in a noncoding region, the [T/C] allelic variation revealed that clones carrying the T allele displayed higher carotenoid accumulation (Figure 4F), suggesting potential effects on regulatory or cis-acting elements. The SNP in *Manes.01G092900* the [G/T] variant also produced a nonsynonymous substitution, replacing methionine with isoleucine. Clones carrying the T allele exhibited higher carotenoid levels (Figure 4F), consistent with the hypothesis that allelic variation at this locus may affect kinase-mediated signaling cascades associated with carotenoid biosynthesis (Ohmura et al., 2001). On chromosome 10, the SNP within Manes.10G117500 resulted in a synonymous change. Although this substitution does not alter the encoded amino acid, individuals carrying the C allele consistently showed higher carotenoid content (Figure. 4F). This pattern suggests that regulatory or structural factors, rather than coding variation, may influence trait expression at this locus.

For dry matter content, *Manes.01G231200* carried a nonsynonymous substitution [T/C] that replaces leucine with lysine, and individuals with the C allele exhibited higher DMC values (Figure 5D). Lysine-bearing variants have been associated with altered protein structural interactions, particularly in cell-wall modification processes that contribute to root developmental dynamics (Yang et al., 2020). On the other hand, *Manes.10G055500* has a synonymous mutation [T/C] affecting Leucine and the reference allele (T) was associated with increased dry matter content (Figure 5E).

Across the genome, favorable allelic combinations were organized into distinct haplotypes. For carotenoids, the haplotypes Hap 9 [TTCC], Hap 11 [ATCC], and Hap 12 [AGCC] exhibited superior accumulation. For dry matter content, the haplotype Hap 3 [CT], combining favorable alleles from both identified loci, showed the most advantageous performance. These results indicate that multi-locus haplotypes capturing combinations of favorable alleles provide more reliable markers for trait improvement than individual SNPs. It is essential to note that certain SNPs (single nucleotide polymorphisms) have a low minor allele frequency (MAF) in the population. Even in the case of superior haplotypes, rare and low-frequency variants contribute significantly to genetic variance. Some studies indicate that non-synonymous variants tend to have lower MAF, and these kinds of variants can have important positive or negative effects on trait expression (Floriani & Lipka, 2025). Including these variants in genomic prediction enhances the reliability of the predictions. This is a crucial consideration because rare alleles are often associated with significant phenotypic variation. If the model misinterprets these alleles and their effects as not statistically significant, it could lose valuable information for genomic prediction. Therefore, it is essential to validate these SNPs in either the training or validation populations in future studies. The identification of these haplotypes represents an important step toward deploying haplotype-based selection strategies in cassava breeding. Because these allelic combinations integrate information across tightly linked loci, they offer improved predictive accuracy for both marker-assisted selection and genomic selection. In addition, these superior haplotypes can guide the selection of parental genotypes for hybridization and backcrossing, facilitating the introgression of favorable alleles into future breeding populations. Incorporating haplotype-level information may therefore accelerate genetic gain for carotenoid biofortification and improved dry matter content in cassava.

### Functional validation and future research directions

GWAS are now widely recognized as an effective approach for dissecting complex quantitative traits in plants, particularly when complemented by haplotype-based analyses. Haplotype information captures historical recombination patterns, including insertions, deletions, and copy number variation, thereby improving the resolution of genomic signals and enhancing the predictive ability of genomic selection models (Briollais et al., 2016; Hamazaki & Iwata, 2020). This makes haplotype-based discovery especially valuable for breeding programs that rely on tagging and tracking favorable genetic variants to accelerate the development of elite cultivars. In this study, integrating GWAS with haplotype analysis using a large, multi-year dataset from the Embrapa breeding program enabled the identification of six candidate genes and their associated SNPs for carotenoid accumulation and dry matter content in cassava. The characterization of favorable haplotypes, specifically [TTCC], [ATCC], [AGCC] for carotenoids and [CT] for dry matter, provides a set of high-value genomic targets that can enhance the accuracy of GS models and inform strategic allelic pyramiding. Haplotype-based genomic selection represents a significant advancement in plant breeding, enhancing the prediction of complex traits by utilizing haplotype blocks. This approach could be a promising strategy for the cassava breeding program in Brazil. The next steps to implement this method include developing Kompetitive Allele-Specific PCR (KASP SNP) markers for the candidate SNPs and validating these markers in independent populations. Once validated, these SNPs could serve as fixed predictors in the next genomic selection cycle.

Using the haplotype blocks to build G_SNP and G_HAP matrices for Bayesian or weighted GBLUP models. In previous research, Lin et al.,(2024) applied haplotype blocks in genomic selection models for yield traits in maize and found that using haplotypes improved prediction accuracy for (GS). Similarly, Difabachew et al., (2023) implemented models using haplotype blocks for 11 traits in winter wheat and explained the advantages of using haplotype blocks instead of individual SNPs, highlighting how this strategy can enhance prediction accuracy in this cultivar. Furthermore, Yu et al., (2006) utilized three LD blocks with KASP molecular markers. The incorporation of these superior haplotypes enabled the breeding program to address low-temperature stress in the new genomic selection population derived from haplotype-based GS. Translating these findings into operational breeding can be facilitated through the development of allele-specific, high-throughput SNP assays, enabling rapid and cost-efficient screening of segregating families. Incorporating these markers into selection pipelines may significantly accelerate genetic gain for carotenoid and dry matter content.

These discoveries also open opportunities for deeper functional investigation. Experimental validation of the identified candidate genes will be essential to confirm their roles in trait expression and to elucidate how specific allelic variants influence transcript and protein behavior. Genome editing technologies such as CRISPR–Cas9 systems offer a compelling toolset for generating targeted knockouts, allelic replacements, or overexpression lines to validate causality and characterize genotype–phenotype relationships (Lin & Musunuru, 2018).

Post-GWAS analyses utilizing omics data represent a compelling approach for future research aimed at accurately identifying causal variants associated with complex traits. By integrating multidimensional datasets, such as transcriptome-wide association studies (TWAS), researchers are able to provide evidence of the relationship between quantitative trait locus (QTL) expression and phenotypic outcomes. Additionally, metabolome (mGWAS) and proteome-wide association studies (PWAS) present further valuable opportunities for exploration. These methodologies yield a more comprehensive understanding of the molecular networks that underpin these traits, highlighting how protein products interact within superior genotypes to enhance carotenoid levels and improve dry matter production. Research focusing on gene-metabolite-trait associations has been pursued by many scientists studying cassava and other crops. For instance, Ding et al., (2023) combined metabolic and phenotypic genome-wide association studies to investigate the genetic architecture of metabolites produced in cassava storage roots. Luo et al., (2023) employed mGWAS and multi-omic functional analyses to explore the mechanisms involved in anthocyanin biosynthesis in cassava leaves. Siriwan et al., (2023) used proteomic and metabolomic analyses to elucidate the mechanisms of disease resistance to mosaic virus disease, uncovering nine proteins and metabolites associated with plant resistance. Parallel transcriptomic and metabolomic analyses of edited or contrasting haplotype carriers would provide further insight into the regulatory networks and biochemical pathways governing carotenoid accumulation and dry matter deposition. Such integrative functional genomics approaches will be crucial to refine gene targets, verify biological mechanisms, and optimize the deployment of favorable alleles within cassava improvement programs.

## Conclusion

This study provides a comprehensive characterization of the genetic architecture underlying carotenoid accumulation and dry matter content in cassava by integrating GWAS with haplotype analysis using an extensive panel of clones from the Embrapa breeding program. This integrative approach led to the identification of six trait-associated SNPs and six biologically compelling candidate genes. Collectively, these loci account for a large proportion of the phenotypic variance—75.56 % for carotenoid content and 76.23 % for dry matter content—demonstrating the high genetic control and clear marker–trait relationships present within the Brazilian germplasm and highlighting their strong potential as robust targets for marker development and genomic-enabled breeding.

The annotated candidate genes encode products central to the biochemical pathways governing these traits. For carotenoids, the implicated genes participate in ABA-related signaling, hydrolase activity essential for lycopene–β-carotene conversion, and fatty-acid biosynthesis, which collectively support plastid development and carotenoid deposition. For dry matter content, the candidate genes are associated with cell wall modification, a process linked to starch accessibility and storage, and glycolytic activity, which contributes to root swelling and biomass accumulation. These functional roles are consistent with the GWAS signals, and the haplotype-phenotype patterns observed across the population.

The combined use of GWAS and haplotype-based analysis enabled the identification of four superior haplotypes carrying favorable alleles for carotenoids and one optimal haplotype for dry matter. These haplotypes represent valuable genetic assets for cassava improvement, providing breeders with high-confidence, trait-linked markers that can be integrated into genomic selection and marker-assisted selection pipelines. Their deployment is expected to enhance prediction accuracy, accelerate allele pyramiding, and increase the efficiency of selecting high-carotenoid and high-dry-matter genotypes. Overall, the results provide an empirical framework for the development of biofortified and high-quality cassava cultivars. They also establish a foundation for future functional studies aimed at validating the roles of the identified genes and clarifying the molecular mechanisms that drive superior performance in specific haplotype classes. Future research will prioritize the development of marker assays utilizing KASP SNPs technology, followed by validation in independent populations. Furthermore, it is essential to perform biological validation of these traits through advanced technologies such as CRISPR-Cas9, transcriptomics, metabolomics, and proteomics. This comprehensive approach will enhance our understanding of the interplay between genes and their protein products, leading to a more profound insight into carotenoid and dry matter production in elite genotypes. This work therefore contributes both actionable breeding tools and new genetic insights that will support long-term genetic gains in cassava improvement programs.

