## Supplementary figures and images for "Uncovering genomic regions controlling root quality traits in Cassava (Manihot esculenta Crantz) using different GWAS models"

### Supplementary_Figure_S1

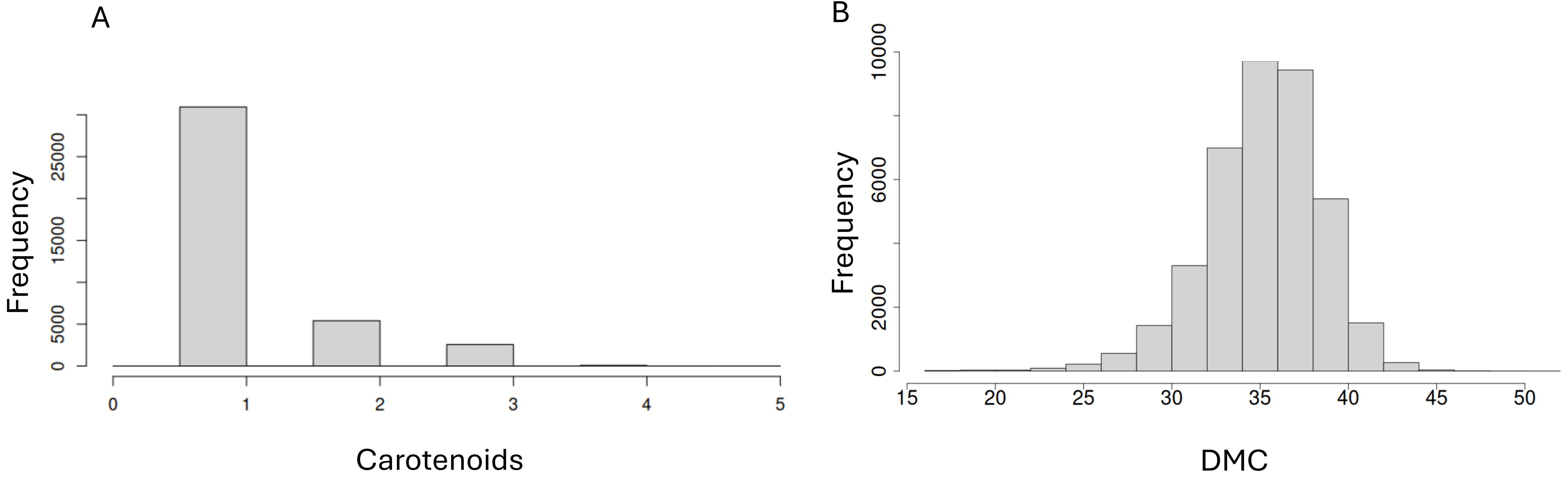

### Supplementary_Figure_S2

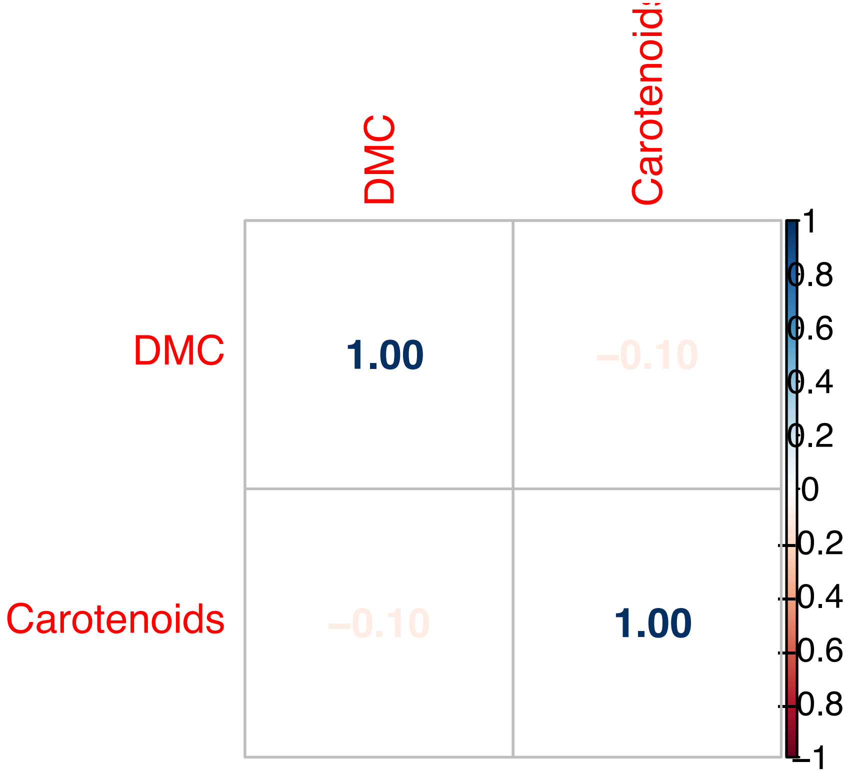

### Supplementary_Figure_S3

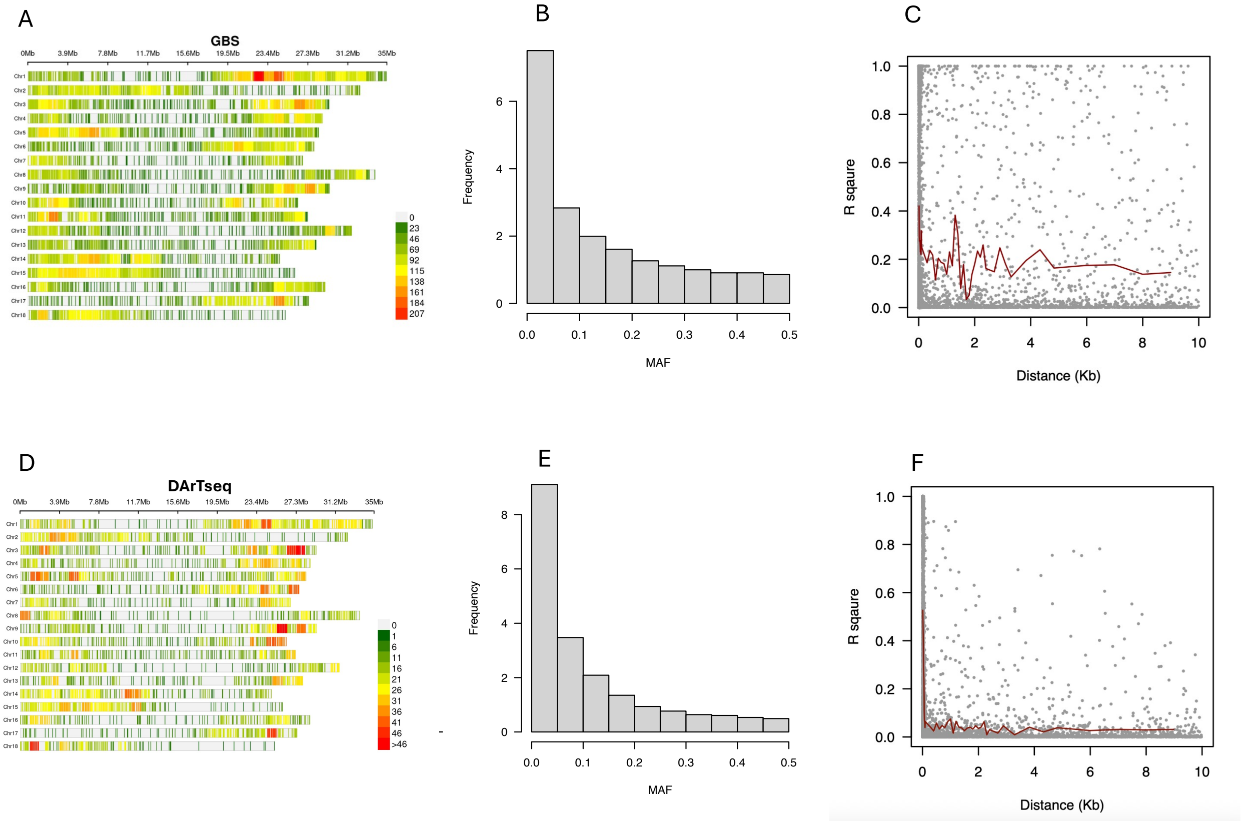
